# Adolescent oligodendrogenesis and myelination restrict experience-dependent neuronal plasticity in adult visual cortex

**DOI:** 10.1101/2023.09.29.560231

**Authors:** Wendy Xin, Megumi Kaneko, Richard H. Roth, Albert Zhang, Sonia Nocera, Jun B. Ding, Michael P. Stryker, Jonah R. Chan

**Affiliations:** Department of Neurology, Weill Institute for Neurosciences, University of California San Francisco; Department of Physiology, Kavli Institute for Fundamental Neuroscience and Weill Institute for Neurosciences, University of California San Francisco; Departments of Neurosurgery and Neurology and Neurological Science, Stanford University

## Abstract

**BACKGROUND:** Developmental myelination is a protracted process in the mammalian brain. One theory for why oligodendrocytes mature so slowly posits that myelination may stabilize neuronal circuits and temper neuronal plasticity as animals age. We tested this hypothesis in the visual cortex, which has a well-defined critical period for experience-dependent neuronal plasticity.

**OBJECTIVES/METHODS:** To prevent myelin progression, we conditionally deleted Myrf, a transcription factor necessary for oligodendrocyte maturation, from oligodendrocyte precursor cells (Myrf cKO) in adolescent mice. To induce experience-dependent plasticity, adult control and Myrf cKO mice were monocularly deprived by eyelid suture. Functional and structural neuronal plasticity in the visual cortex were assessed in vivo by intrinsic signal optical imaging and longitudinal two photon imaging of dendritic spines, respectively.

**RESULTS:** During adolescence, visual experience modulated the rate of oligodendrocyte maturation in visual cortex. Myrf deletion from oligodendrocyte precursors during adolescence led to inhibition of oligodendrocyte maturation and myelination that persisted into adulthood. Following monocular deprivation, visual cortex activity in response to visual stimulation of the deprived eye remained stable in adult control mice, as expected for post-critical period animals. By contrast, visual cortex responses to the deprived eye decreased significantly following monocular deprivation in adult Myrf cKO mice, reminiscent of the plasticity observed in adolescent mice. Furthermore, visual cortex neurons in adult Myrf cKO mice had fewer dendritic spines and a higher level of spine turnover. Finally, monocular deprivation induced spatially coordinated spine size decreases in adult Myrf cKO, but not control, mice.

**CONCLUSIONS:** These results demonstrate a critical role for oligodendrocytes in shaping the maturation and stabilization of cortical circuits and support the concept of myelin acting as a brake on neuronal plasticity during development.

## Introduction

Relative to other cellular developmental processes, oligodendrogenesis and myelination occur in an extremely protracted manner, spanning the first few postnatal weeks in mice and the first three decades in humans. A longstanding theory for explaining this unique timing posits that developmental myelination restricts the ability of cortical neurons to undergo experience-dependent plasticity in adulthood (*1–5*). The progression of myelination within the mouse visual cortex supports this idea, where an accumulation of myelin within the input layer of the cortex coincides with the closure of a critical period for experience-induced functional neuronal plasticity (*1*). Furthermore, sensory deprivation induces myelin remodeling within visual cortex (*6*), suggesting a potential link between sensory experience, myelination, and neuronal function. To better define the relationship between visual experience and oligodendrocyte dynamics, we used genetic lineage tracing to track the generation of mature oligodendrocytes in the adolescent visual cortex following visual deprivation. To directly test the idea that developmental myelination restricts neuronal plasticity, we genetically inhibited the generation of mature oligodendrocytes and progression of myelination during adolescence and assessed functional and structural experience-induced neuronal plasticity in the adult visual cortex. These experiments address a fundamental, as yet untested hypothesis about brain development and inform our broader understanding of how oligodendrocytes and myelin influence neuronal circuit function and plasticity.

### Sensory experience modulates oligodendrocyte lineage dynamics in the adolescent visual cortex

Mature, myelinating oligodendrocytes arise from oligodendrocyte precursor cells (OPCs). By crossing a mouse line that specifically expresses an inducible Cre in OPCs (NG2CreER) (*7*) with a Cre-dependent reporter (Tau-mGFP) (*8*), we can identify newly generated oligodendrocytes by mGFP expression, as only cells that are OPCs at the time of tamoxifen administration will express the reporter. To determine whether sensory experience during adolescence can influence oligodendrocyte maturation, we gave tamoxifen to 4-week-old NG2CreER:Tau-mGFP mice and used eyelid suture to monocularly deprive them for ten days, then quantified oligodendrogenesis in the visual cortex (Figure 1A). OPCs that recombined the reporter were detected by co-expression of mGFP and the OPC-specific protein PDGFRα, whereas new pre-myelinating oligodendrocytes expressed mGFP but not PDGFRα, and new mature oligodendrocytes expressed mGFP and the myelin protein MBP (Figure 1B; S1A). Overall, the contralateral cortex - which primarily receives visual inputs from the deprived eye - contained fewer mGFP^+^ cells than the ipsilateral cortex (Figure 1C, H). By lineage stage, there were slightly fewer mGFP^+^ OPCs, as well as fewer PDGFRα^+^ OPCs overall, in the contralateral cortex (Figure 1D, I; S1B). The percentage of recombined OPCs was similar in both hemispheres (Figure S1C). There was no difference in the number of new pre-myelinating oligodendrocytes between the two hemispheres (Figure 1E, J), though this stage of oligodendrocyte maturation is transient and identified by the absence of immature and mature oligodendrocyte markers (Figure 1B) and, therefore, less well defined. New mature oligodendrocyte density was most affected by monocular deprivation, with significantly fewer cells present in the contralateral cortex (Figure 1F, K).

**Figure 1.**
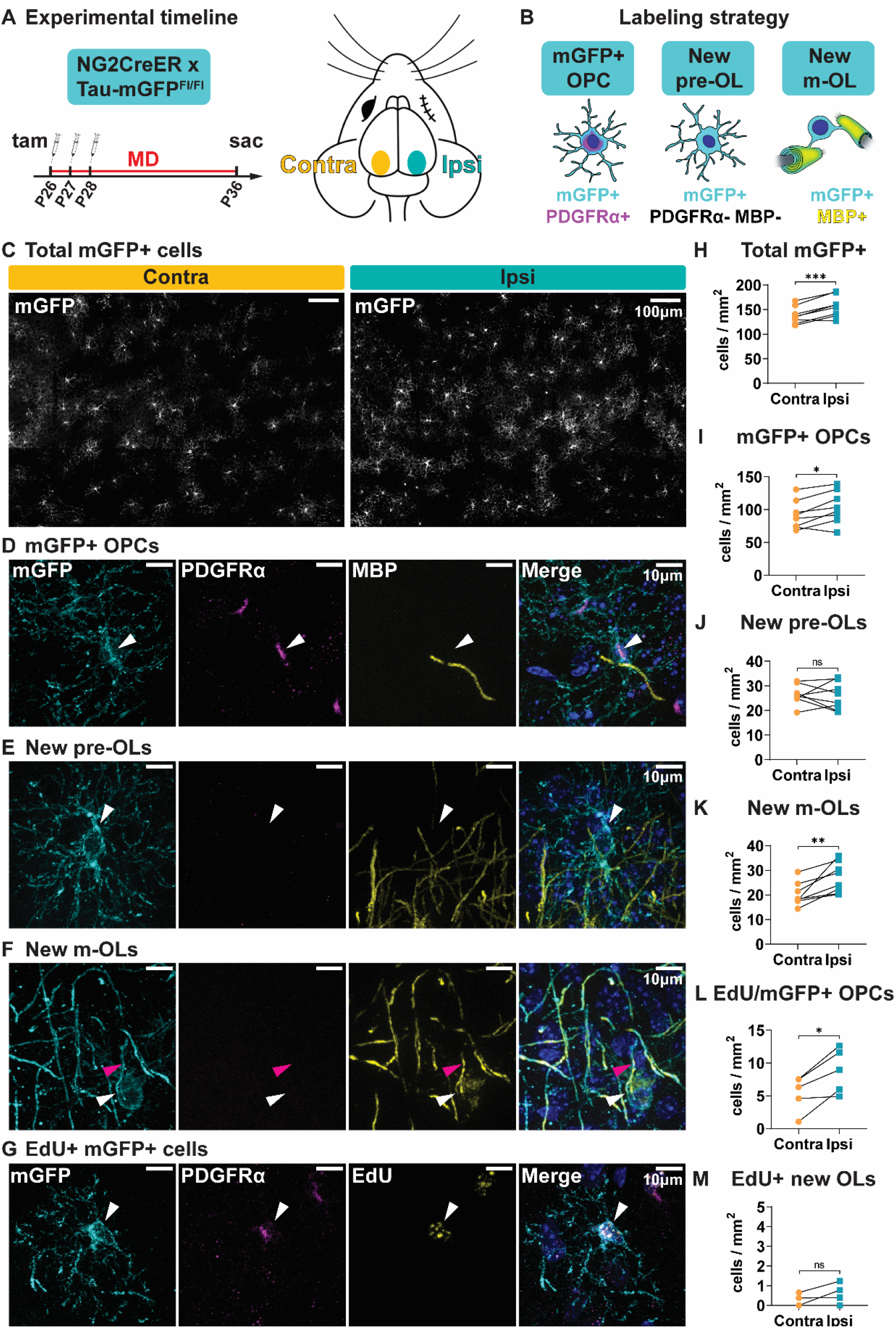
Sensory experience during adolescence modulates oligodendroglial dynamics. (**A**) Experimental strategy and timeline. Tam = tamoxifen; MD = monocular deprivation. (**B**) Labeling strategy for identifying oligodendrocyte precursor cells (OPC), newly formed pre-myelinating oligodendrocytes (pre-OL), and mature oligodendrocytes (m-OL). (**C**) Representative images and (**H**) quantification of mGFP^+^ cells from both hemispheres of visual cortex. Contra = hemisphere contralateral to the deprived eye; ipsi = hemisphere ipsilateral to the deprived eye. (**D**) Example image and (**I**) quantification of mGFP^+^ OPCs. (**E**) Example image and (**J**) quantification of newly formed pre-OLs. (**F**) Example image and (**K**) quantification of newly formed m-OLs. (**G**) Example image and (**L**) quantification of mGFP^+^ EdU^+^ OPCs. (**M**) Quantification of mGFP^+^ EdU^+^ PDGFRα^-^ newly formed OLs. White arrowheads = cell body; pink arrowheads = MBP^+^ mGFP^+^ myelin sheath. Statistical details in Table S1. *p<0.05, **p<0.01, ***p<0.001, ns = not significant.

There are two potential explanations for the decrease in both OPCs and mature oligodendrocytes in the contralateral cortex. Sensory deprivation may modulate OPC proliferation, which changes the density of OPCs and, indirectly, the density of mature oligodendrocytes that would be generated. The other possibility is that sensory deprivation modulates oligodendrocyte differentiation and myelination, in which case OPC proliferation may change due to a decreased drive to replace differentiated cells. To distinguish between these two possibilities, we administered the thymidine analog 5-Ethynyl-2’-deoxyuridine (EdU), which can be incorporated into the newly synthesized DNA of dividing cells, at the same time as tamoxifen (Figure S1D). After ten days of monocular deprivation, there were fewer EdU^+^ cells overall (Figure S1E, F) and fewer EdU^+^ OPCs in the contralateral cortex, indicating fewer OPCs proliferated in the contralateral cortex during the experimental window (Figure 1G, L). However, the number of newly generated oligodendrocytes that arose from proliferated OPCs (mGFP^+^ EdU^+^ PDGFRα^-^) was the same in both hemispheres, which in most mice was between 0-1 cell per mm^2^ (Figure 1M), compared to ∼100 overall newly generated oligodendrocytes (Figure 1I). Thus, the majority of new oligodendrocytes generated during the ten days of monocular deprivation arose from OPCs that did not first proliferate. As such, an indirect effect of altered proliferation cannot account for the difference in mature oligodendrocyte density between contralateral and ipsilateral cortex. Taken together, these results suggest that sensory deprivation during adolescence alters the rate of oligodendrocyte differentiation and myelination in the visual cortex.

### OPC-specific Myrf deletion in adolescent mice prevents oligodendrogenesis and myelination

Having established that sensory experience regulates oligodendrocyte maturation during adolescence, we next asked whether blocking adolescent oligodendrogenesis could alter neuronal function and plasticity within the visual cortex. We generated PDGFRαCreER:Myrf^Fl/Fl^ mice, which enables OPC-specific (*9*) deletion of Myrf, a transcription factor that is necessary for oligodendrocyte differentiation (*10*). To prevent adolescent oligodendrogenesis, we administered tamoxifen to CreER^+^ (cKO) and CreER^-^ (control) mouse pups from P10 to P14 (Figure 2A). In control mice, there was a rapid accumulation of ASPA^+^ mature oligodendrocytes from 4 to 8 weeks of age, whereas the number of oligodendrocytes in cKO mice plateaued at 4 weeks of age (Figure 2B, E; S2). Accordingly, the pattern of myelination in visual cortex remained sparse and patchy in 8-week-old cKO mice (Figure 2D, S2). Given the global nature of the manipulation, we also assessed oligodendrocyte density in earlier stages of the visual pathway, i.e. optic nerve and lateral geniculate nucleus, at 8 weeks. Optic nerve oligodendrocyte density was unchanged, and myelination grossly intact, in Myrf cKO mice (Figure S3A). In the lateral geniculate nucleus, we did observe a decrease in the number of oligodendrocytes in cKO mice, although the decrease was smaller and overall patterns of myelination were less affected than in visual cortex (Figure S2, S3B). The heterogeneous effect of Myrf deletion along the visual pathway is likely driven by differences in the timing of developmental oligodendrogenesis, wherein oligodendrocytes in proximal portions of the visual pathway like optic nerve differentiate much earlier (*11*) and are therefore not affected by subsequent Myrf deletion.

**Figure 2.**
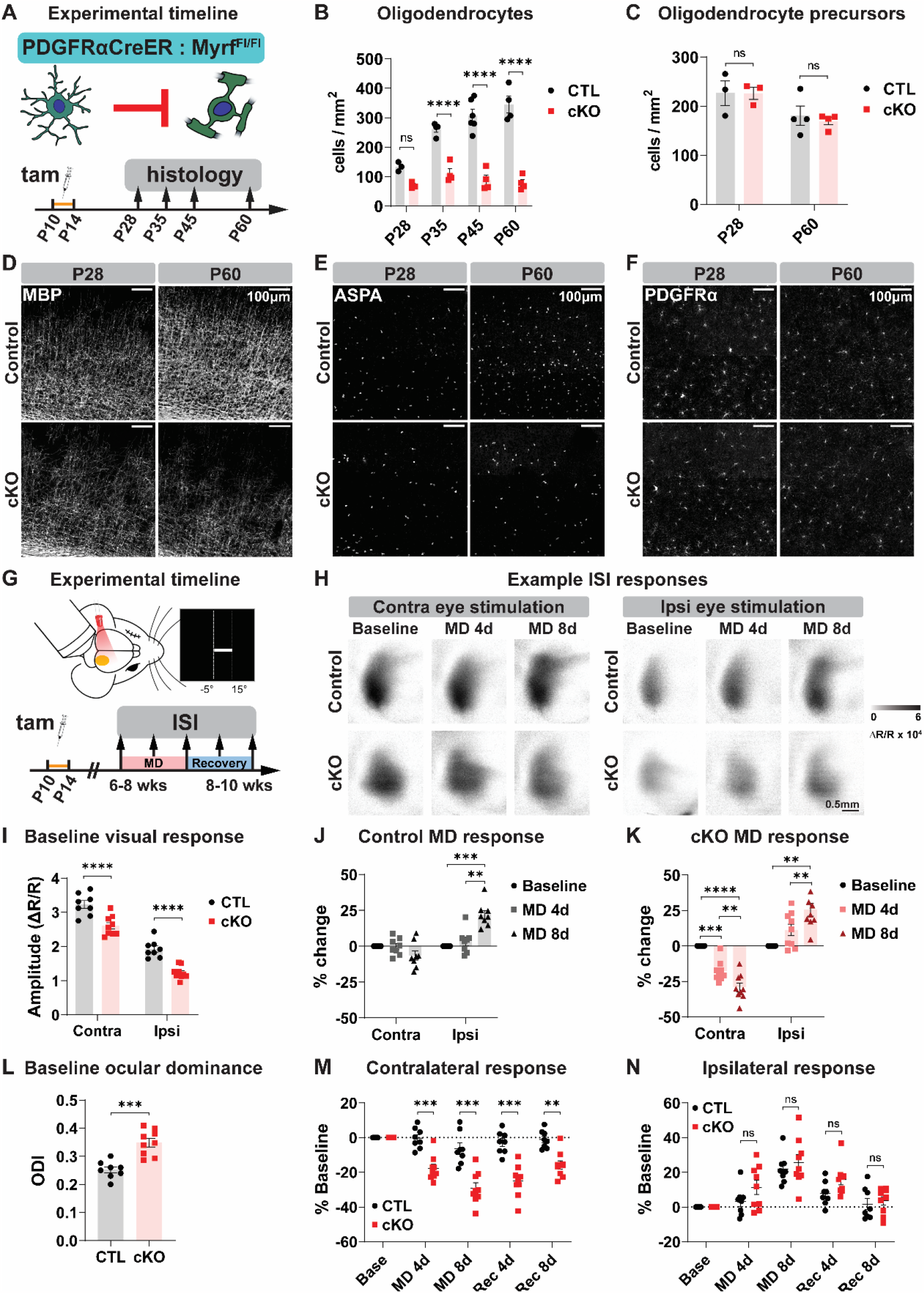
Impairing adolescent oligodendrogenesis disrupts adult visual cortex activity and enhances experience-dependent neuronal plasticity. (**A**) Experimental strategy and timeline for histology. Tam = tamoxifen. (**B**) Quantification and (**E**) example images of ASPA+ mature oligodendrocytes in the visual cortex at P28 and P60 in CreER^-^ (CTL) and CreER^+^ (cKO) mice. (**C**) Quantification and (**F**) example images of PDGFRα+ OPCs in visual cortex. (**D**) Example images of myelin in visual cortex. (**G**) Experimental strategy and timeline for intrinsic signal imaging (ISI). MD = monocular deprivation. (**H**) Example images from ISI. MD 4/8d = after 4 or 8 days of monocular deprivation. (**I**) Amplitude of ISI responses and (**L**) ocular dominance in binocular visual cortex at baseline. Contra = ISI response to contralateral eye stimulation, ipsi = ISI response to ipsilateral eye stimulation. ODI = ocular dominance index, defined as (contra – ipsi ISI responses) / (contra + ipsi). (**J**) Change in binocular visual cortex ISI responses in adult control mice following four (MD4) or eight (MD8) days of MD. (**K**) Change in ISI responses in adult cKO mice. (**M**) Change in contralateral visual cortex ISI responses following MD and recovery (Rec). (**N**) Change in ipsilateral visual cortex ISI responses. Statistical details in Table S1. **p<0.01, ***p<0.001, ****p<0.0001, ns = not significant.

Myrf deletion did not alter the density of OPCs in adolescent or adult visual cortex (Figure 2C, F), likely due to the exquisite ability of OPCs to maintain homeostatic density in vivo (*12*). However, blocking differentiation could push OPCs to continuously proliferate instead, with homeostatic density being maintained by a simultaneous increase in cell death. To investigate this possibility, we injected adult Myrf cKO and littermate control mice with EdU for five days to assess the rate of OPC proliferation in visual cortex. Similar to our observations with sensory deprivation, decreasing differentiation – in this case by genetic manipulation – was associated with a decrease in OPC proliferation (Figure S4). We also did not detect any increased cell death in adult visual cortex (Figure S5A), overt gliosis as indicated by GFAP immunoreactivity (Figure S5B), or any changes in the density of astrocytes (Figure S5C) or microglia (Figure S5D). Thus, Myrf deletion potently inhibits oligodendrogenesis and myelination in visual cortical areas without grossly altering the density and morphology of other glial populations.

### Adolescent oligodendrogenesis limits visual cortex responsiveness to visual stimulation and experience-dependent visual cortex plasticity in adulthood

Myelination has long been hypothesized to be important for regulating cortical neuronal maturation and plasticity (*2, 5, 13, 14*), but the functional consequence of preventing developmental oligodendrogenesis and myelination has not been directly tested. We first verified that the loss of adolescent oligodendrogenesis does not induce neurodegeneration in visual cortex by immunostaining with a neurofilament light chain (NF-L) antibody that binds an epitope of NF-L accessible only in degenerating axons (*15*). As a positive control, we detected axonal degeneration in the spinal cord of mice that underwent experimental autoimmune encephalitis, particularly in regions with myelin loss (Figure S6A). By contrast, we saw little to no degeneration in the adult visual cortex of control and Myrf cKO mice (Figure S6B). Thus, unlike demyelination, a lack of developmental myelination does not result in axonal degeneration.

To assess functional neuronal activity in the visual cortex, we used intrinsic signal optical imaging, which enables non-invasive longitudinal monitoring of bulk neuronal activity during visual stimulation (*16*). Adult mice were lightly anesthetized and head fixed, and a visual stimulus was delivered to the ipsilateral or contralateral eye in the binocular portion of the mouse visual field (Figure 2G). Myrf cKO mice exhibited normal retinotopic organization in the visual cortex (Figure S7A), as expected, given that cortical retinotopy is established by P15 (*17*) and oligodendrocyte density remains comparable between control and Myrf cKO mice before P28 (Figure 2B, E). However, visual cortex responses to visual stimulation of either eye were significantly weaker in Myrf cKO mice compared to control mice, with a more pronounced decrease in ipsilateral responsiveness (Figure 2I). As a result, baseline ocular dominance, i.e. the relative strength of visual cortex responses to stimulation of the contralateral vs. ipsilateral eye, was significantly higher in Myrf cKO mice (Figure 2L). These results indicate that adolescent oligodendrogenesis and myelination are required for proper visual cortex maturation and function in adulthood.

It has previously been proposed that myelin in the visual cortex may gate the ability of neurons to undergo experience-dependent plasticity (*1, 2, 4, 13*). However, no study to date has functionally tested this hypothesis by cell type-specific manipulation of oligodendrocyte maturation or myelination. Therefore, we examined experience-dependent plasticity in the adult visual cortex of control and Myrf cKO mice. A hallmark of critical period experience-dependent plasticity is the loss of visual cortex responsiveness to the deprived eye following a brief period of monocular deprivation (*18–20*). In wildtype adult mice, monocular deprivation induces an increase in visual cortex response to the non-deprived eye, but no change in response to the deprived eye (*18, 20*). Indeed, in adult control mice, we observed stable visual cortex responses to the deprived eye after four or eight days of monocular deprivation, though visual cortex responses to the non-deprived eye increased as expected (Figure 2H, J; S7B). By contrast, adult Myrf cKO mice exhibited pronounced decreases in visual cortex responsiveness to the deprived eye after monocular deprivation (Figure 2H, K; S7C). Comparing control and Myrf cKO visual cortex responses, we found that contralateral responses after monocular deprivation were significantly different between groups (Figure 2M), whereas ipsilateral responses were comparable (Figure 2N). These results demonstrate a key role for adolescent oligodendrogenesis and myelin in restricting functional experience-dependent neuronal plasticity in the visual cortex.

### Adolescent oligodendrogenesis regulates structural synaptic plasticity in adult visual cortex

The decrease in visual cortex plasticity from critical period to adulthood has been linked to several cellular factors. Most notably, disrupting parvalbumin interneuron maturation or perineuronal net formation in the visual cortex results in increased visual cortex plasticity (*21–25*). We did not observe any gross differences in the density of parvalbumin neurons, nor in the presence of perineuronal nets, within the visual cortex of Myrf cKO and control mice (Figure S8). Another anatomical correlate for functional experience-dependent neuronal plasticity is the physical rewiring of visual cortex circuitry (*26–30*). In adolescent mice, monocular deprivation induces the elimination of dendritic spines in visual cortex pyramidal neurons (*26, 27, 29*). To test whether adolescent oligodendrogenesis and myelination can also regulate structural neuronal plasticity, we crossed Myrf cKO mice to Thy1-YFP-H mice (*31*), which allowed us to visualize a subset of pyramidal neurons and their dendritic spines in the visual cortex. We performed in vivo two photon longitudinal imaging in adult mice to track dendritc spines in the contralateral cortex over a period of normal vision and monocular deprivation (Figure 3A, B). Overall, spine density was lower in Myrf cKO mice than littermate control mice (Figure 3C, D), echoing the decrease in functional visual cortex responses we observed in cKO mice (Figure 2I). The relative decrease in spine density was maintained throughout the period of monocular deprivation (Figure 3D). Spine turnover, on the other hand, was higher in Myrf cKO mice (Figure 3E), which was driven by an increase in spine eliminations (Figure 3F) as well as a trend towards increased spine additions (Figure 3G).

**Figure 3.**
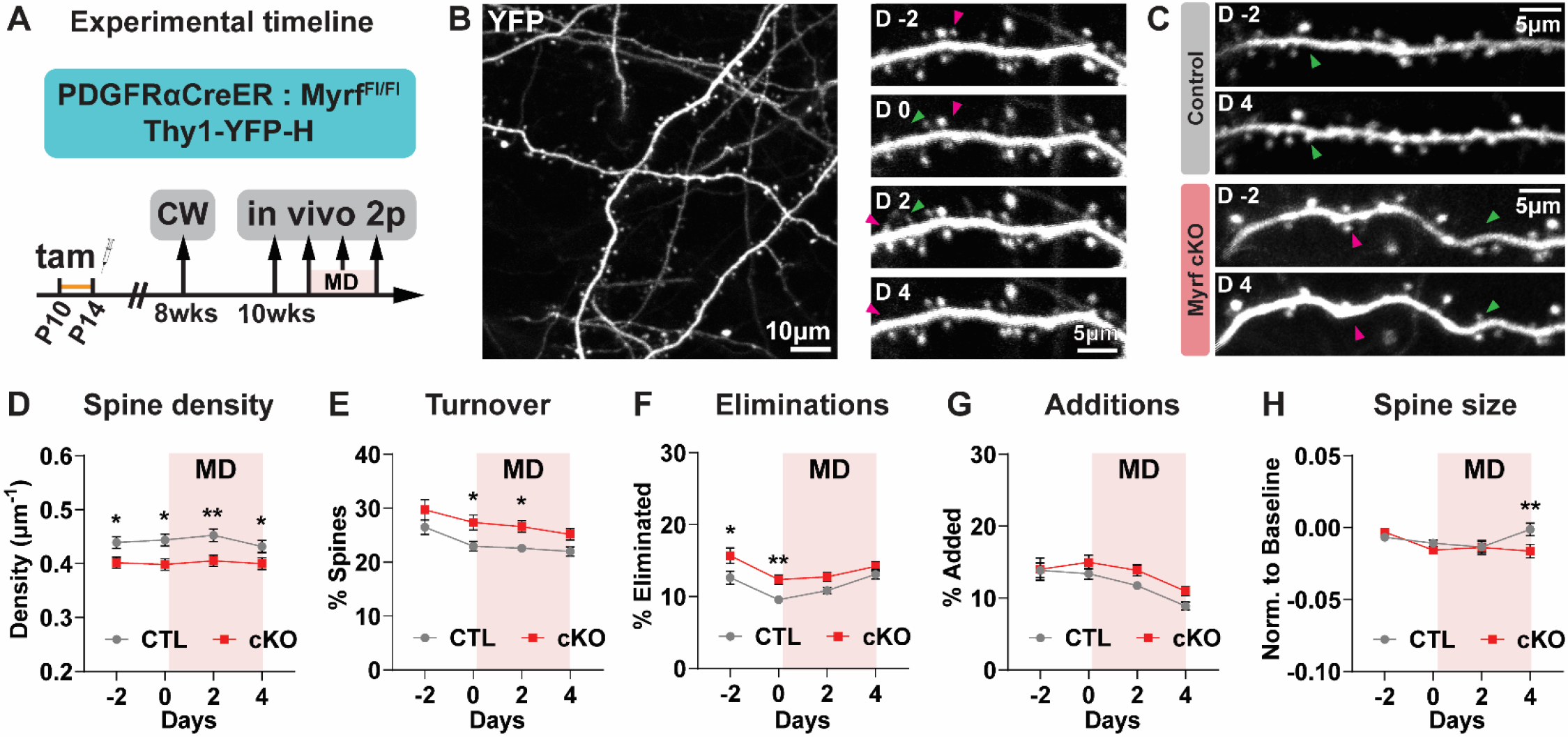
Adult mice with impaired adolescent oligodendrogenesis have fewer spines and higher spine turnover. (**A**) Experimental strategy and timeline. Tam = tamoxifen; CW = cranial window surgery; MD = monocular deprivation. (**B**) Left: example 2-photon image from an adult mouse visual cortex, contralateral to the deprived eye. Right: Example dendrite imaged at 2 days pre-MD (D-2), just prior to MD (D 0), two days following MD (D 2), and four days following MD (D 4). (**C**) Example dendrites from control and cKO mice. (**D**) Average spine density, (**E**) spine turnover, (**F**) spine eliminations (**G**) spine additions, and (**H**) spine size in control (CTL) and cKO visual cortex dendrites. Green arrowheads = spine that was added; pink arrowheads = spine that was eliminated. Data presented as mean +/- SEM. Statistical details in Table S1. *p<0.05, **p<0.01.

Spines that were added or eliminated during the imaging sessions accounted for approximately one third of the total population (Figure 3E), meaning most spines remained present throughout the experiment. However, persistent spines can still exhibit functionally significant structural plasticity; indeed, spine size is tightly correlated with synaptic strength (*32, 33*). Overall, we observed a difference in spine size changes after four days of monocular deprivation, with cKO mice exhibiting a decrease in relative spine size compared to control mice (Figure 3H), as well as a negative (leftwards) shift in the distribution of spine size changes (Figure S9A), meaning more spines decreased in size with monocular deprivation. For both groups, the size change of each spine after two days of monocular deprivation was strongly associated with its size change after four days of monocular deprivation, meaning spines that increased or decreased after two days continued to increase or decrease after four days (Figure S9B, C).

Previous studies examining experience-induced spine plasticity have found that experience-induced spine plasticity tends to be spatially clustered along segments of dendrites (*34, 35*). This spatial clustering is critical for the functional consequences of spine plasticity and leads to supra-linear summation, both electrically and biochemically, in the dendritic integration of individual changes in synaptic strength (*36–39*). To test whether spine size changes were spatially clustered following adult monocular deprivation, we performed a nearest neighbor analysis, where the directional change of each spine was compared to the directional change of its nearest spine neighbor (Figure 4A). In control mice, there was no correlation between the direction of spine size change between nearest neighbors (Figure 4B; S9D). By contrast, there was a significant positive correlation between the size changes of nearest spine neighbors in cKO mice (Figure 4B; S9E). At the level of dendrites, the percentage of spine pairs that increased together was similar between groups (Figure 4C), but cKO dendrites had a higher percentage of spine pairs that decreased together (Figure 4D). Combined, cKO dendrites had more spine pairs changing in the same direction than in control dendrites (Figure 4E), and a similar percentage of spine pairs changing in the opposite direction (Figure 4F). These results indicate that spine changes induced by monocular deprivation are more spatially clustered in cKO mice.

**Figure 4.**
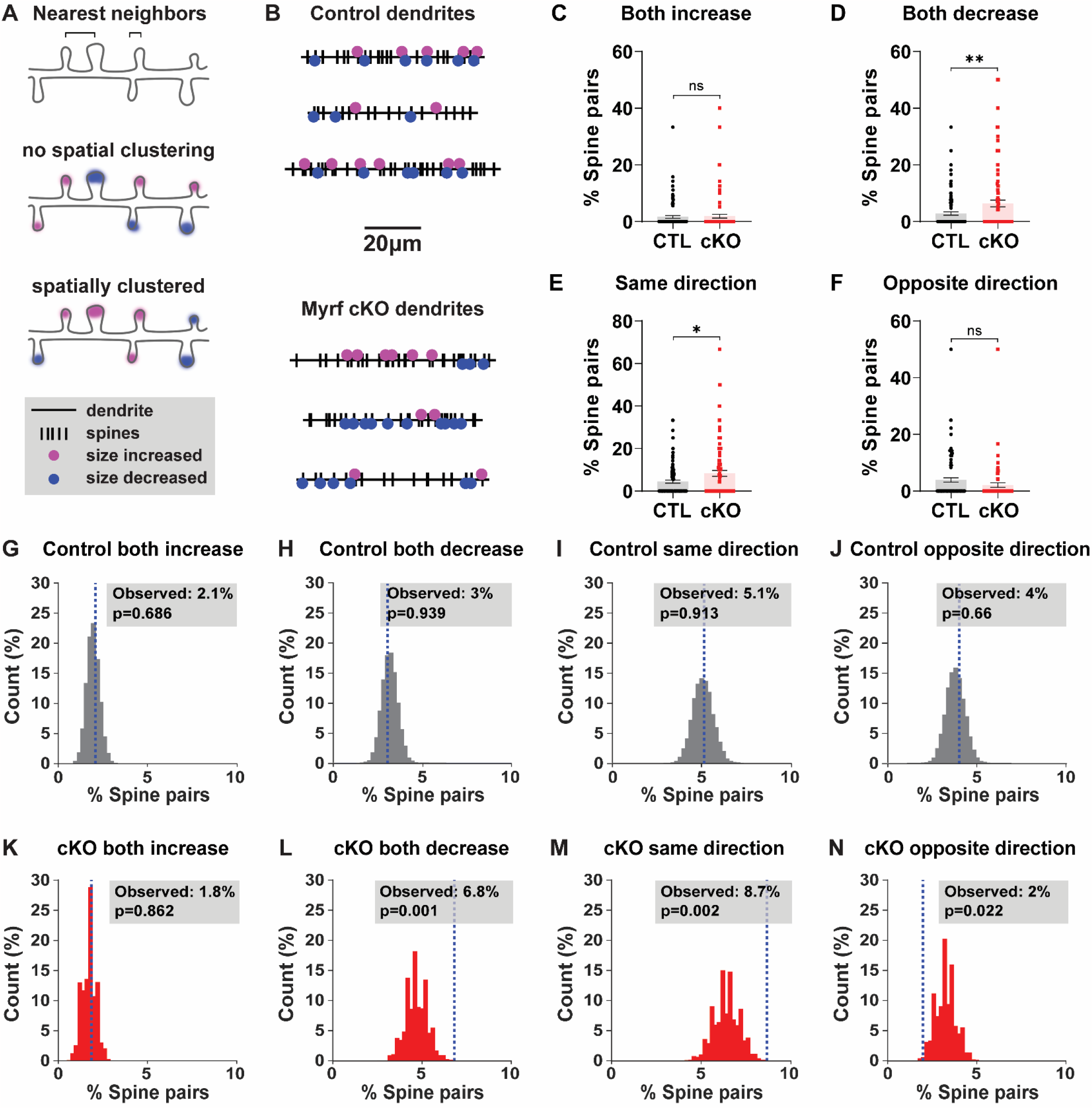
Monocular deprivation induces spatially clustered spine size decreases in adult mice with impaired adolescent oligodendrogenesis. (**A**) Schematic of nearest neighbor analysis. (**B**) Spinodendrograms of three example dendrites per group. (**C**) Percentage of spine pairs per dendrite that both increased, decreased (**D**), changed in the same direction (**E**), or changed in opposite directions (**F**) in control (CTL) or cKO mice. (**G**) Distribution of percentage of spine pairs that increased together from 10000 random spine/location pairings in control mice; blue dashed line marks the observed percentage of spine pairs. (**H**) Distribution of percentages of random spine pairs that decreased together, changed in the same direction (**I**), or changed in opposite directions (**J**) in control mice. (**K**) Distribution of percentages of random spine pairs (and observed percentage, blue dashed line) that increased, decreased (**L**), changed in the same direction (**M**), or changed in opposite directions (**N**) in cKO mice. Statistical details in Table S1. *p<0.05, **p<0.01, ns = not significant.

Given that we observed a greater number of spines decreasing in size in cKO mice (Figure 3H; S9A), it is possible that the higher level of spine change clustering in cKO mice is simply due to more spine decreases present along each dendrite. To determine whether this was the case, we compared the observed spine pairs with a distribution of spine pairs generated by randomly shuffling the changes in spine size along all spine positions in each dendrite (Figure S9F, G). In control mice, the observed spine pairs increasing or decreasing together fell within the distribution of random spine pairs, indicating that the level of spine clustering we observed was occurring at chance levels (Figure 4GJ). In cKO mice, the percentage of spine pairs increasing together fell within the distribution of random spine pairs (Figure 4K), but the percentage of spine pairs decreasing together was significantly higher than would be predicted by chance (Figure 4L). Altogether, we observed many more spine pairs changing in the same direction (Figure 4M), and fewer spine pairs changing in the opposite direction (Figure 4N), in cKO mice than expected based on the overall number of spines increasing and decreasing in size. Thus, cKO mice exhibit an enhanced level of spatially clustered decreases in spine size following monocular deprivation. These differences in structural plasticity likely underlie the selective decrease in functional visual cortex responses we observed following monocular deprivation in adult cKO mice.

## Discussion

A role for oligodendrocytes and myelination in regulating neuronal circuit structure and function across development has long been suspected (*1–4, 14*). Here, using cell type-specific genetic approaches, we find that sensory deprivation during adolescence reduces oligodendrocyte maturation and that blocking adolescent oligodendrogenesis enhances neuronal plasticity in the adult visual cortex. Our results provide crucial empirical evidence for the longstanding theory that developmental myelination restricts experience-dependent neuronal plasticity and identify a novel non-neuronal mechanism for regulating coordinated changes in dendritic spine plasticity. Although our study focuses on the role of developmental myelination, the ability for oligodendrocytes and myelin to modulate neuronal function may expand beyond development. Recent studies have reported a requirement for adult oligodendrogenesis in multiple forms of learning and memory (*40–43*), but it is still unclear how oligodendrogenesis or myelin plasticity is modulating the underlying circuit activity to influence behavior (*40, 44*). Based on our findings within a sensory circuit, it is plausible that new oligodendrocytes and myelin may serve to stabilize the formation of new synaptic connections following learning to enable long-term memory. This possibility is further supported by imaging studies in vivo reporting neuronal cell type-specific and activity driven changes in myelin sheaths following sensory manipulations or motor learning (*6, 45*). Our results demonstrate the potential for oligodendrocytes and myelin to act as a toggle between neuronal circuit stability and reorganization for lifelong brain plasticity and implicate myelin as a key driver of circuit dysfunction in neurodevelopmental disorders presenting with impaired oligodendrocyte maturation (*40, 46, 47*).

## Methods

### Animals

All mice were handled in accordance with, and all procedures approved by, the Institutional Animal Care and Use Committee of the University of California San Francisco. Mice were group housed and given food and water ad libitum on a 12h light/dark cycle. Males and females were used for all experiments. To track oligodendrogenesis during adolescence, NG2CreER:tau-mGFP mice (Zhu et al. 2011, Jax # 008538; Hippenmeyer et al. 2005, Jax # 021162) received 100 mg/kg tamoxifen (Sigma cat# T5648) by oral gavage from P26-28. A subset also received 80 mg/kg 5-Ethynyl-2’-deoxyuridine (Carbosynth cat# NE08701) by intraperitoneal injection at the same time. To block adolescent oligodendrogenesis, PDGFRαCreER:MyrfFl/Fl mice (Kang et al. 2010, Jax # 018280; Emery et al. 2009, Jax # 010607) received 100 mg/kg tamoxifen by oral gavage from P10-14. To visualize dendritic spines in vivo, PDGFRαCreER:MyrfFl/Fl mice were crossed to Thy1-YFP-H mice (Jax # 003782).

### Immunohistochemistry

Mice were deeply anesthetized with Avertin and perfused transcardially with 4% paraformaldehyde in 1X PBS. Brain tissue was isolated and postfixed in this solution overnight at 4°C, then stored in 1X PBS with 0.1% NaAz. Brains were sucrose protected (30% in PBS) prior to flash freezing and sectioned coronally (30 μm) on a sliding microtome. Free-floating sections were permeabilized/blocked with 0.2% Triton X-100 and 10% normal goat serum in 1X PBS for 1hr at room temperature. Sections were incubated with primary antibodies prepared in 0.2% Triton X-100 and 10% normal goat serum in 1X PBS at 4°C overnight. Sections were incubated with secondary antibodies in 10% normal goat serum in 1X PBS for 2hr at room temperature. Primary antibodies and concentrations used are listed in Table S2. Primary antibodies have been validated for use in immunohistochemistry in mouse tissue, in published literature, and on the manufacturer’s websites. Secondary antibodies used included the following: Alexa Fluor 488−, 594−, or 647−conjugated secondary antibodies to rabbit, mouse, goat, chicken, rat or guinea pig (1:1000; all raised in goat; Jackson ImmunoResearch). Cell nuclei were labeled with DAPI (Vector Laboratories).

### Fixed tissue imaging and analysis

Tiled z stacks (with 2 μm steps) spanning 30 μm sections of visual cortex and lateral geniculate nucleus, or 20 μm sections of optic nerve, were taken with a Zeiss Axio Imager Z1 with ApoTome attachment and Axiovision software, using a 10X objective. For quantification, images were taken from 2-3 sections per mouse. Cell density was quantified manually using Cell Counter in Fiji. Experimenters were blinded to genotype through imaging acquisition and analysis.

### Monocular deprivation

Mice were anesthetized using 5% isofluorane, and anesthesia was maintained with 2-3% isofluorane. The right eyelid was sutured close by two mattress stitches either at P26 for adolescent NG2CreER:tau-mGFP mice or between 6-12 weeks for post-critical period PDGFRαCreER:MyrfFl/Fl mice. Meloxicam and buprenorphine were administered pre- and post-surgery for pain management. Animals were checked daily to ensure the sutured eye remained closed for the required duration of the experiment. Sutures were removed just prior to post-monocular deprivation intrinsic signal imaging sessions. Eyes were flushed with sterile saline and checked for clarity under a microscope. Only mice without corneal opacities or signs of infection were used.

### Intrinsic signal imaging

Repeated optical imaging of intrinsic signals and quantification of ocular dominance were performed as described (*16*). Briefly, during recording mice were anesthetized with 0.7% isoflurane in oxygen applied via a homemade nose mask, supplemented with a single intramuscular injection of 20 to 25 μg chlorprothixene. Mice underwent a non-invasive procedure where a headplate was fixed to the surface of the skull to enable head-fixed imaging, and images were recorded transcranially. Intrinsic signal images were obtained with a Dalsa 1M30 CCD camera (Dalsa, Waterloo, Canada) with a 135 × 50 mm tandem lens (Nikon Inc.) and red interference filter (610 ± 10 nm). Frames were acquired at a rate of 30 fps, temporally binned by 4 frames, and stored as 512 × 512 pixel images after binning the 1024 × 1024 camera pixels by 2 × 2 pixels spatially. The visual stimulus for recording the binocular zone, presented on a 40 × 30 cm monitor placed 25 cm in front of the mouse, consisted of 2°-wide bars, which were presented between −5° and 15° on the stimulus monitor (0° = center of the monitor aligned to center of the mouse) and moved continuously and periodically upward or downward at a speed of 10°/s. The phase and amplitude of cortical responses at the stimulus frequency were extracted by Fourier analysis as described (1). Response amplitude was an average of at least 4 measurements. Ocular dominance index (ODI) was computed as described (*16*). Briefly, the binocularly responsive region of interest (ROI) was chosen based on the ipsilateral eye response map after smoothing by low-pass filtering using a uniform kernel of 5 × 5 pixels and thresholding at 40% of peak response amplitude. OD score (C−I)/(C+I) was computed for each pixel in this ROI, where C and I represent the magnitude of response to contralateral and ipsilateral eye stimulation, followed by calculation of the ODI as the average of OD score for all responsive pixels. Experimenters were blinded to genotype through imaging and analysis.

### Cranial window surgery

At the age of 8-12 weeks a square 3x3 mm cranial window (#1 coverslip glass, Warner Instruments) was placed over the left hemisphere of the cortex, contralateral to the deprived eye. Mice were anesthetized using 5% isofluorane, and anesthesia was maintained with 2-3% isofluorane. A craniotomy matching the size of the coverslip was cut using #11 scalpel blades (Fine Science Tools) and the coverslip was carefully placed on top of the dura within the craniotomy without excessive compression of the brain. The window was centered using stereotactic coordinates 2 mm lateral and 3 mm posterior from bregma for visual cortex. The window and skull were sealed using dental cement (C&B Metabond, Parkell). A custom-made metal head bar was attached to the skull during surgery for head-fixed imaging. Mice were allowed to recover for 2-3 weeks before two-photon imaging.

### In vivo longitudinal imaging

Longitudinal in vivo two-photon imaging was performed as previously described with PDGFRαCreER:MyrfFl/Fl mice crossed to Thy1-YFP-H mice. Specifically, apical dendrites of L5 pyramidal neurons were imaged repeatedly 10-100 μm below the cortical surface through the cranial window in mice under isoflurane anesthesia. Images were acquired using a Bergamo II two-photon microscope system with a resonant scanner (Thorlabs) and a 16 x / 0.8 NA water immersion objective lens (Nikon). YFP was excited at 925 nm with a mode-locked tunable ultrafast laser (InSightX3, Spectra-Physics) with 15-100 mW of power delivered to the back-aperture of the objective. Image stacks were acquired at 1,024 × 1,024 pixels with a voxel size of 0.12 μm in x and y and a z-step of 1 μm. Imaging frames from resonant scanning were averaged during acquisition to achieve a pixel dwell time equivalent of 1 ns. Up to six imaging regions were acquired for each mouse. Representative images shown in figures were created by making z-projections of 3D stacks and were median filtered and contrast enhanced.

### Analysis of in vivo spine imaging

Dendritic spines were analyzed using the custom software Map Manager (https://mapmanager.net) written in Igor Pro (WaveMetrics) as previously described (*35, 48*). Experimenters were blinded to genotype through imaging acquisition and analysis. For annotation, the dendritic shaft was first traced using a modified version of the “Simple Neurite Tracer” plugin in ImageJ. Spine positions along a dendritic segment were manually identified by the location of the spine tip in 3D image stacks of all imaging sessions. For longitudinal analysis, spines were further tracked across days by comparing images from different sessions and connecting persistent spines. Spine addition and elimination rates are calculated as number of newly added or eliminated spines on a given imaging session divided by the total number of spines of that dendritic segment on the previous imaging session. The turnover ratio represents the sum of spine addition and elimination.

The fluorescence intensity of dendritic spines was used as a proxy for spine size. Therefore, a three-dimensional ROI was defined for each spine, the dendritic shaft (4 μm stretch) adjacent to that spine, and a nearby background region. To compare intensity values between imaging sessions and account for small variations in daily imaging conditions, the spine signal intensity was normalized to the signal on the adjacent dendritic shaft after background subtraction. Each spine value was subsequently normalized to an average of the baseline imaging sessions by first subtracting the baseline value and then dividing over the sum of the baseline and respective imaging day value. This normalizes spine size change values between -1 and 1. All spine analysis was performed for each dendritic segment, averaged per genotype, and is presented as the average of values from two adjacent imaging sessions (−3 and -2, -1 and 0, 1 and 2, etc.) to increase clarity.

For analysis of spine clustering, spines were classified as increasing, decreasing, or stable based on their average change in size on days 1-4 compared to baseline. The threshold for these categories was set based on the variability in control mice and was defined at baseline ± one standard deviation of size changes (±0.14). Nearest neighbor analysis was calculated by finding the closest neighbor of every spine along each dendritic segment. Each nearest neighbor pair was only included once in the dataset and pairs were excluded if their distance was below 1.0 μm (to avoid overlapping ROIs) or above 3.5 μm. The fraction of nearest neighbor spine pairs in which both spine increase, both decrease, changes occur in the same direction, or in opposite directions, were quantified to compare the degree of clustering between genotype.

To test the statistical significance of clustering, the nearest neighbor analysis was performed on a pool of randomized spines in which the spine size change values were randomly shuffled along all spine positions in each dendrite. A Monte-Carlo p-value was calculated by summing the tail of the histograms from 10,000 pools of randomized spine pairings in which the nearest neighbor analysis resulted in spine pair fractions that exceeded the real observed value.

### Statistical analysis

All graphed values are shown as mean ± SEM. Statistical details of experiments (statistical tests used, statistical values, exact n) are listed in Table S1. The number of animals included in each experiment was based on standards established in the literature. Statistics were performed using GraphPad Prism. Statistical significance was defined as having a p value of less than 0.05. Tests for normality and equal variances were used to determine the appropriate statistical test to employ. All reported t tests were two-tailed, with Welch’s correction when group variances were significantly different. For experiments with more than two groups, one-way ANOVAs were used. For experiments with more than one variable, two-way ANOVAs were used. For experiments with repeated-measurements from the same animals, two-way repeated-measures ANOVAs were used.

## Acknowledgements

We are grateful to all members of the Chan lab for their insightful feedback on the manuscript.

## Authors contributions

W.X. and J.R.C. conceived the experiments. W.X., M.K., R.H.R., A.Z., and S.N. performed experiments. W.X., M.K., and R.H.R. analyzed data. J.B.D., M.P.S., and J.R.C. provided resources and scientific guidance. W.X. wrote the manuscript with input from all authors.

## Data and materials availability

All data are available in the main text or the supplementary materials. Request for materials and raw data should be addressed to W.X. and J.R.C.

## Supplementary files

Figs. S1 to S9

Tables S1 and S2

