## Supplementary Files for "Adolescent oligodendrogenesis and myelination restrict experience-dependent neuronal plasticity in adult visual cortex"

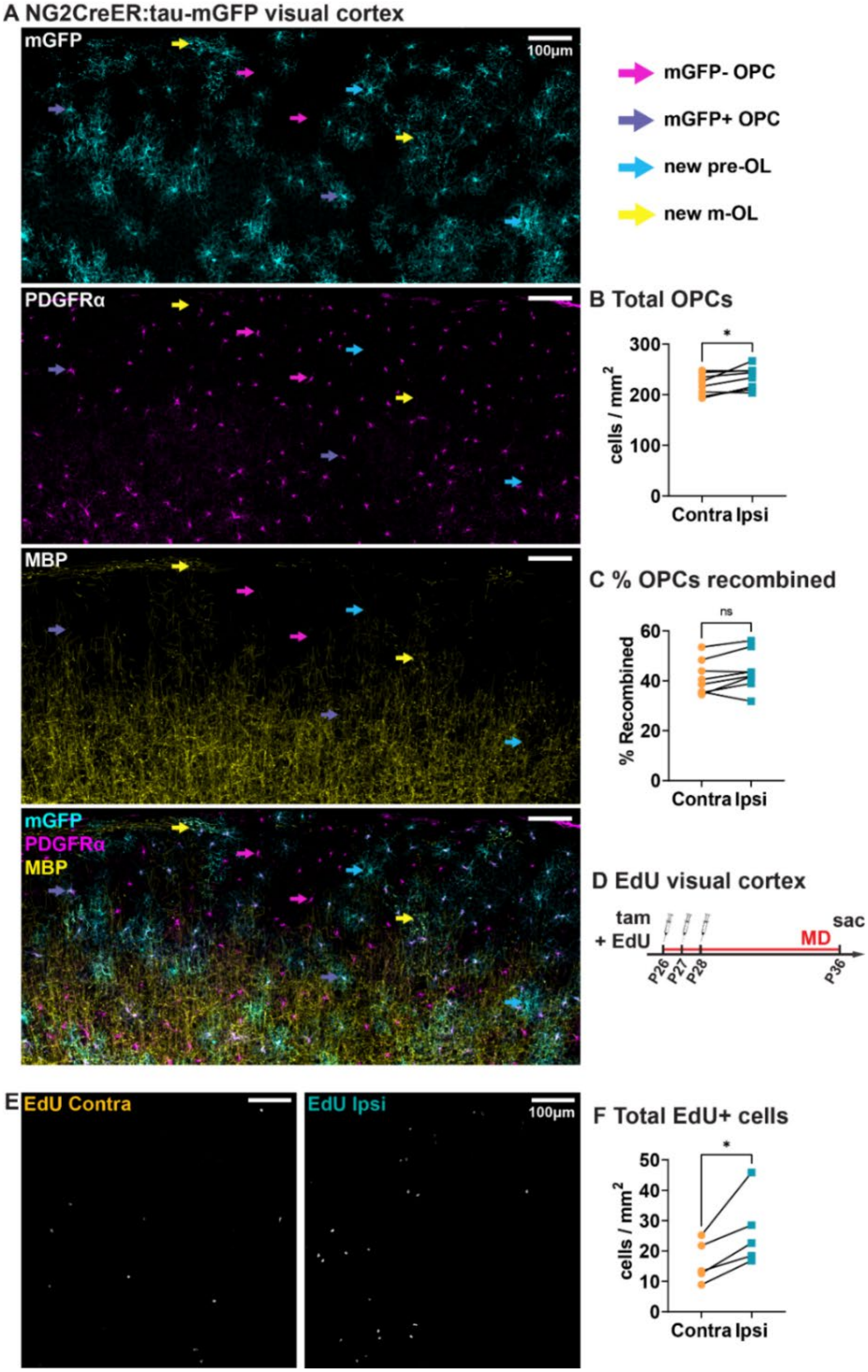


**Fig. S1.**

**Tracking adolescent oligodendroglial dynamics using NG2CreER:tau-mGFP mice.** (**A**) Example image from visual cortex. OPC = oligodendrocyte precursor cell, pre-OL = pre-myelinating oligodendrocyte, m-OL = mature oligodendrocyte. (**B**) Total PDGFRα$^{+}$ OPCs per hemisphere. Contra = hemisphere contralateral to the deprived eye, ipsi = hemisphere ipsilateral to the deprived eye. (**C**) Percentage of PDGFRα$^{+}$ OPCs that were mGFP$^{+}$. (**D**) Timeline of EdU injections. (**E**, **F**) Example images and quantification of total EdU+ cells in both hemispheres of visual cortex. Statistical details in Table S1. *p<0.05, ns = not significant.

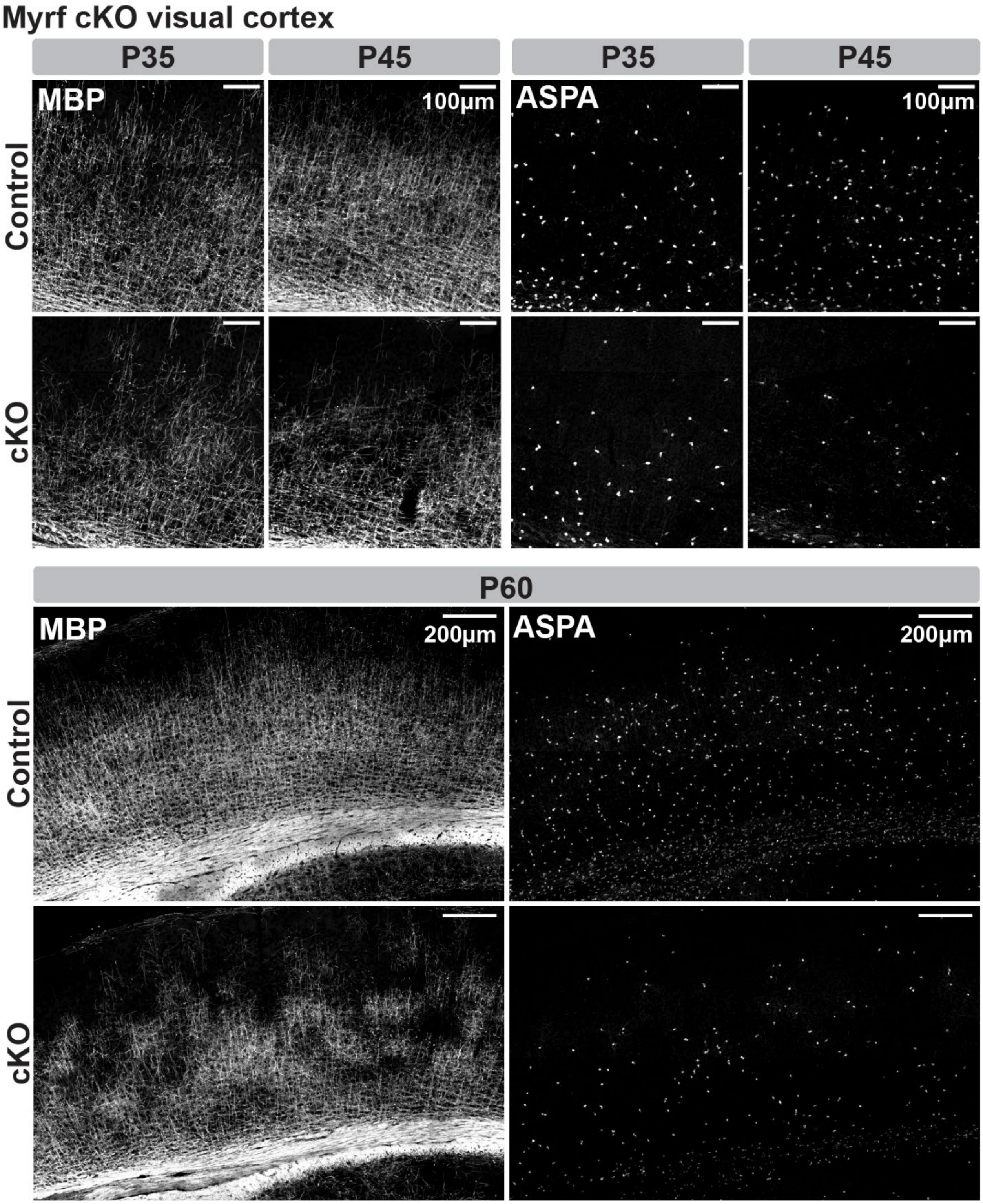


**Fig. S2.**

**OPC-specific deletion of Myrf in adolescence impairs oligodendrogenesis and myelination in visual cortex**. Example images of myelin (MBP) and mature oligodendrocytes (ASPA) in visual cortex of control and Myrf cKO mice at P35, P45, and P60.

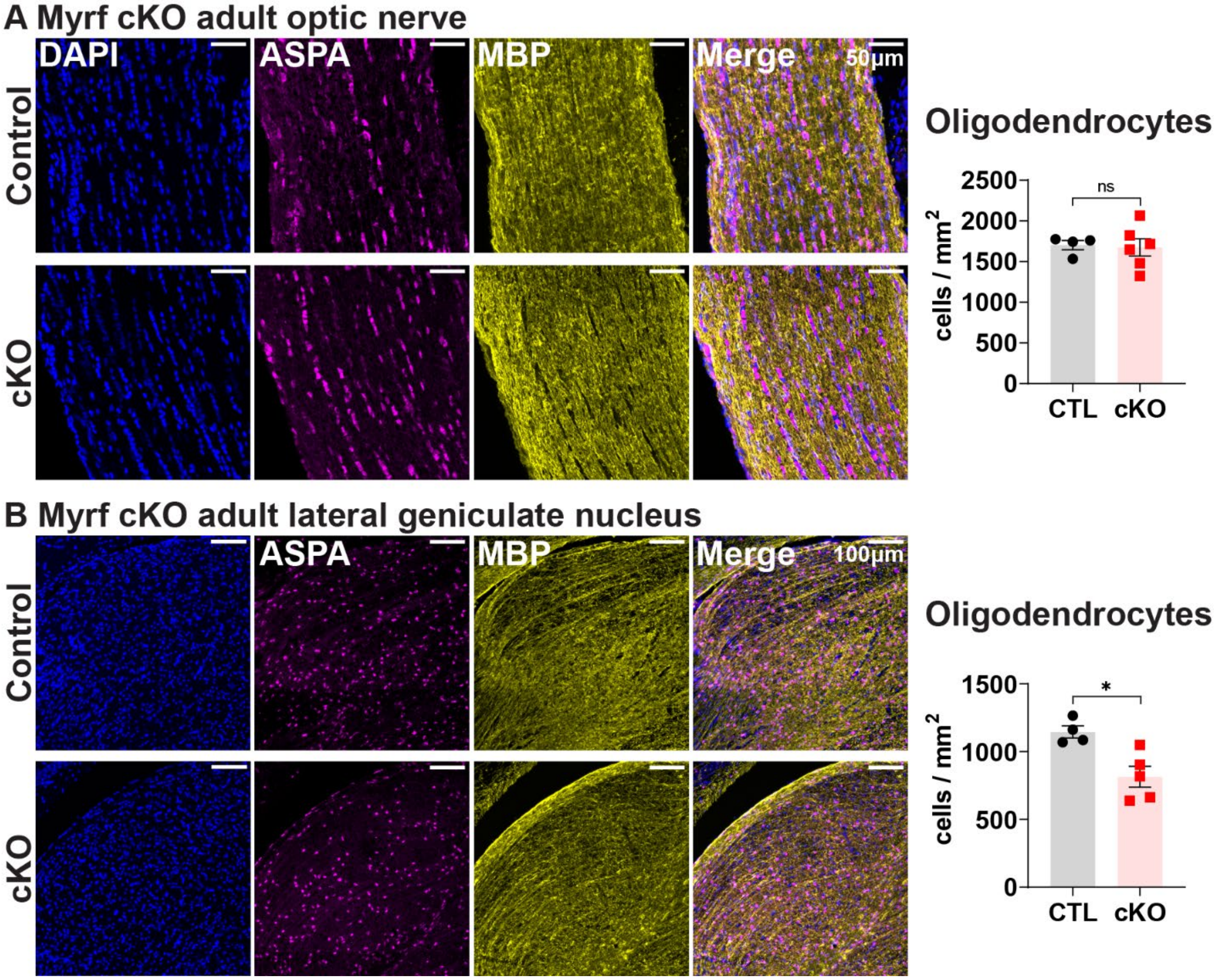


**Fig. S3.**

**Oligodendrogenesis and myelination in the optic nerve and lateral geniculate nucleus of adult control and Myrf cKO mice.** (**A**) Example images of myelin (MBP) and example images/quantification of mature oligodendrocytes (ASPA) in optic nerve. (**B**) Example images of myelin and and example images/quantification of mature oligodendrocytes in lateral geniculate nucleus. Statistical details in Table S1. *p<0.05, ns = not significant.

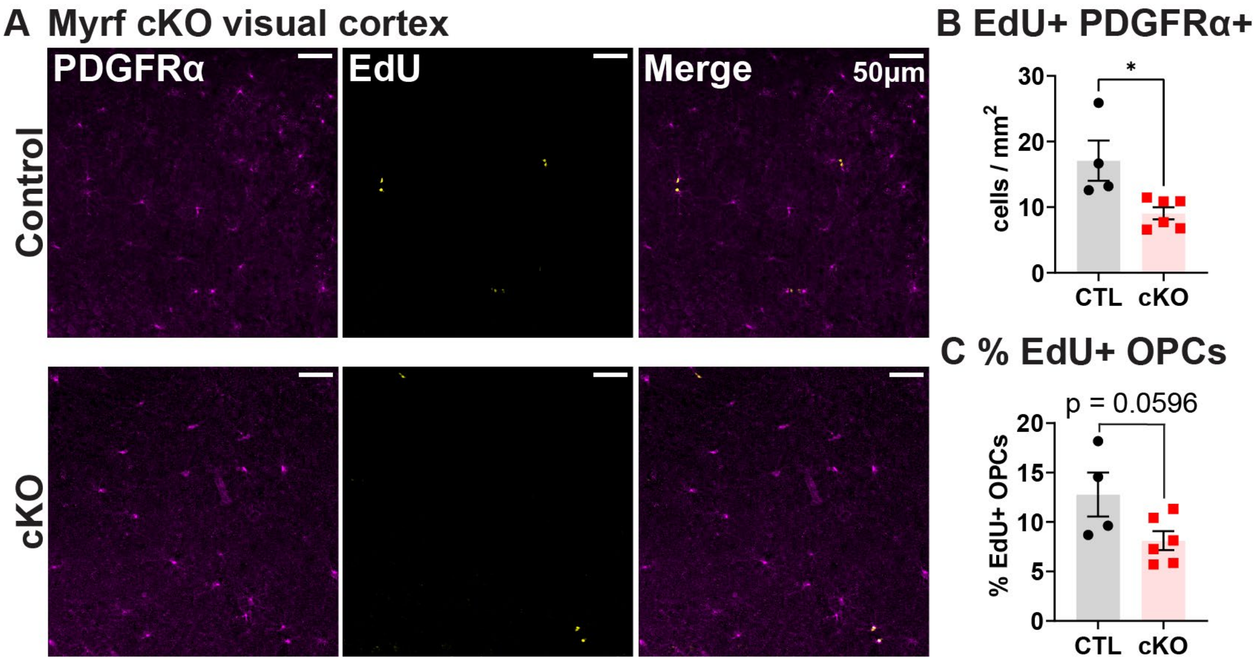


**Fig. S4.**

**OPC proliferation in visual cortex of adult control and Myrf cKO mice.** (**A**) Example images of OPCs (PDGFRα) and EdU labeling in visual cortex. (**B**) Quantification of PDGFRα$^+$ EdU$^+$ OPCs in visual cortex. (**C**) Quantification of percentage of PDGFRα$^+$ OPCs that were EdU$^+$ in visual cortex. Statistical details in Table S1. *p<0.05.

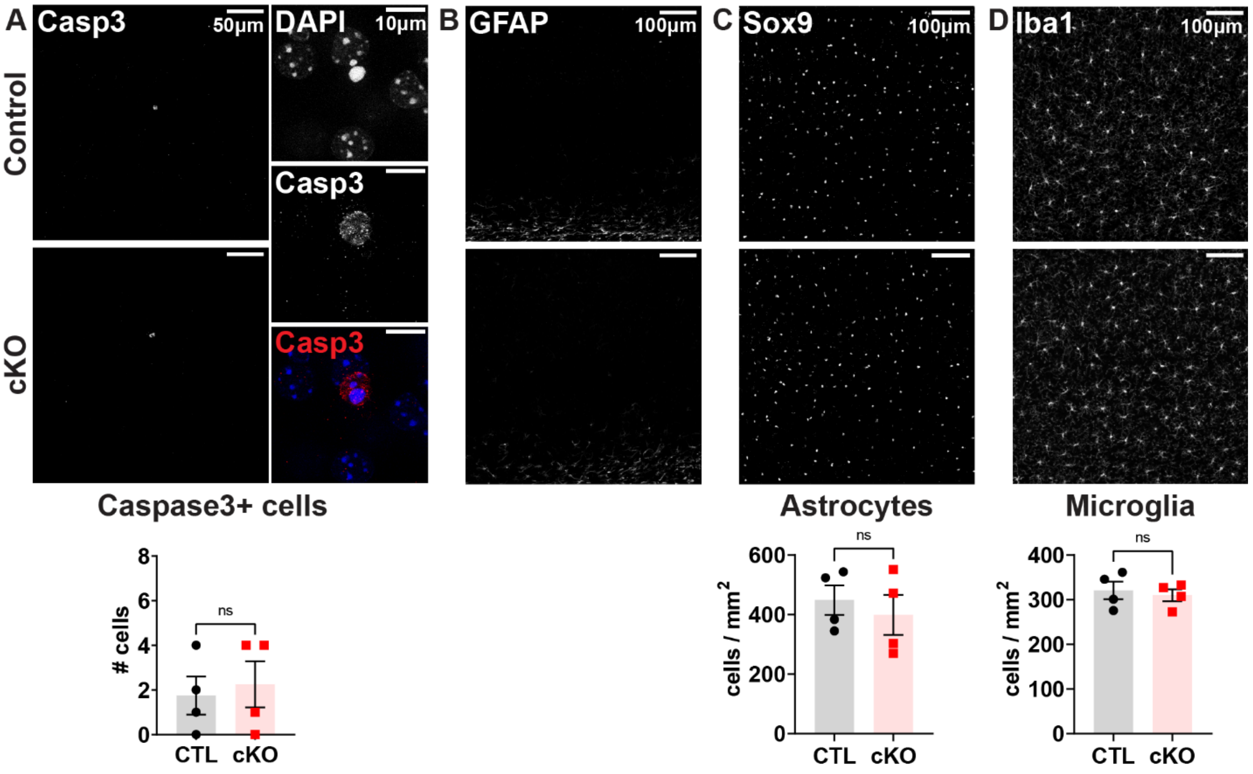


**Fig. S5.**

**Cell death, astrocyte density, and microglia density in adult control and Myrf cKO mice.** (**A**) Example images and quantification of Caspase$^{+}$ cells in visual cortex. (**B**) Example images of GFAP immunostaining in visual cortex. (**C**) Example images and quantification of Sox9$^{+}$ astrocytes in visual cortex. (**D**) Example images and quantification of Iba1$^{+}$ microglia in visual cortex. Statistical details in Table S1. ns = not significant.

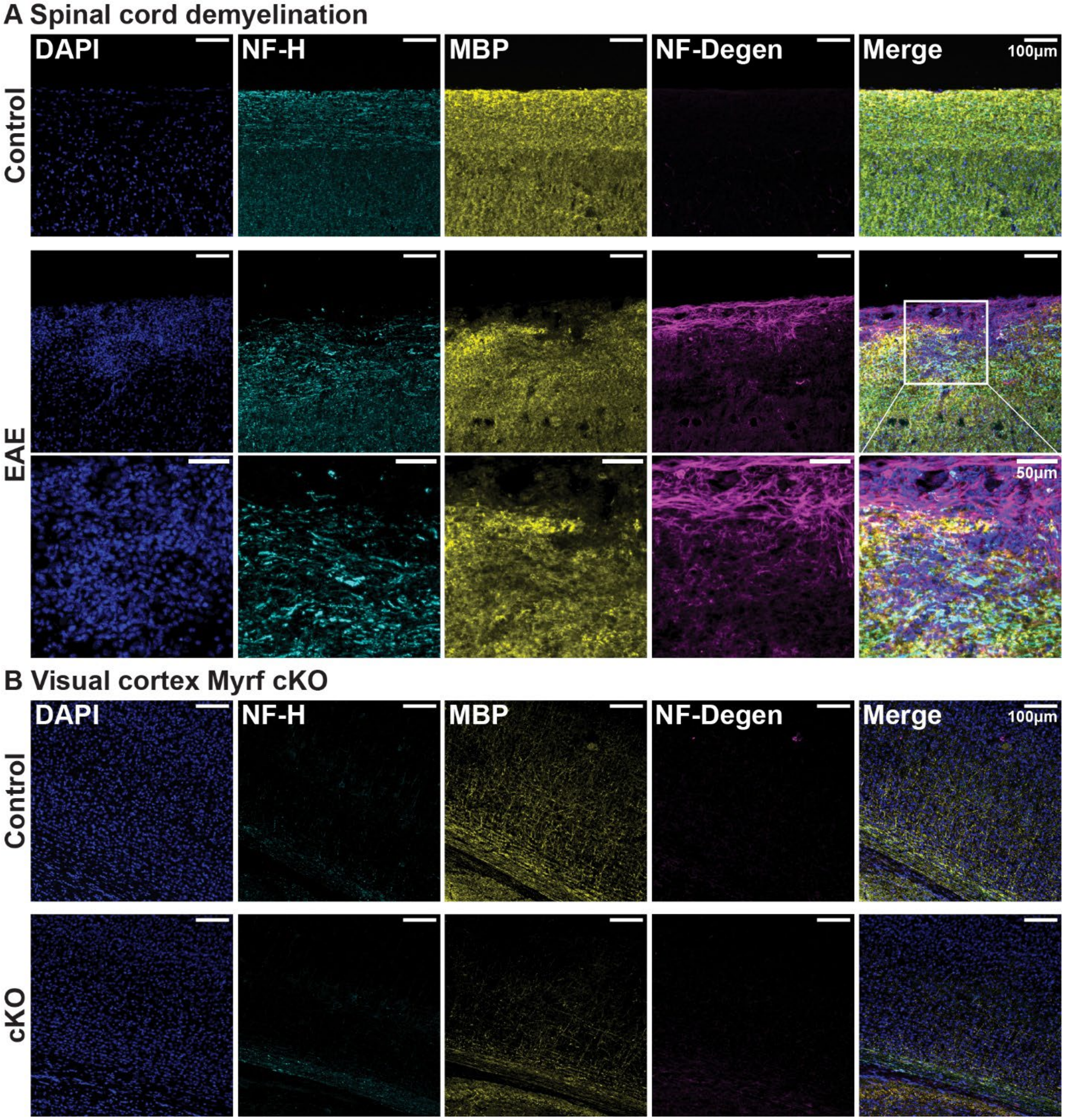


**Fig. S6.**

**Assessing neurodegeneration in demyelination and in Myrf cKO mice.** (**A**) Immunostaining for neurofilament H (NF-H), myelin basic protein (MBP), and neurofilament light chain DegenoTag (NF-Degen) in spinal cords of control mice and mice that underwent experimental autoimmune encephalitis (EAE). Disordered NF-H and prominent NF-Degen signal can be detected in the spinal cord of EAE mice, most notably in regions of demyelination. (**B**) Immunostaining for NF-H, MBP, and NF-Degen in visual cortex of control and Myrf cKO mice.

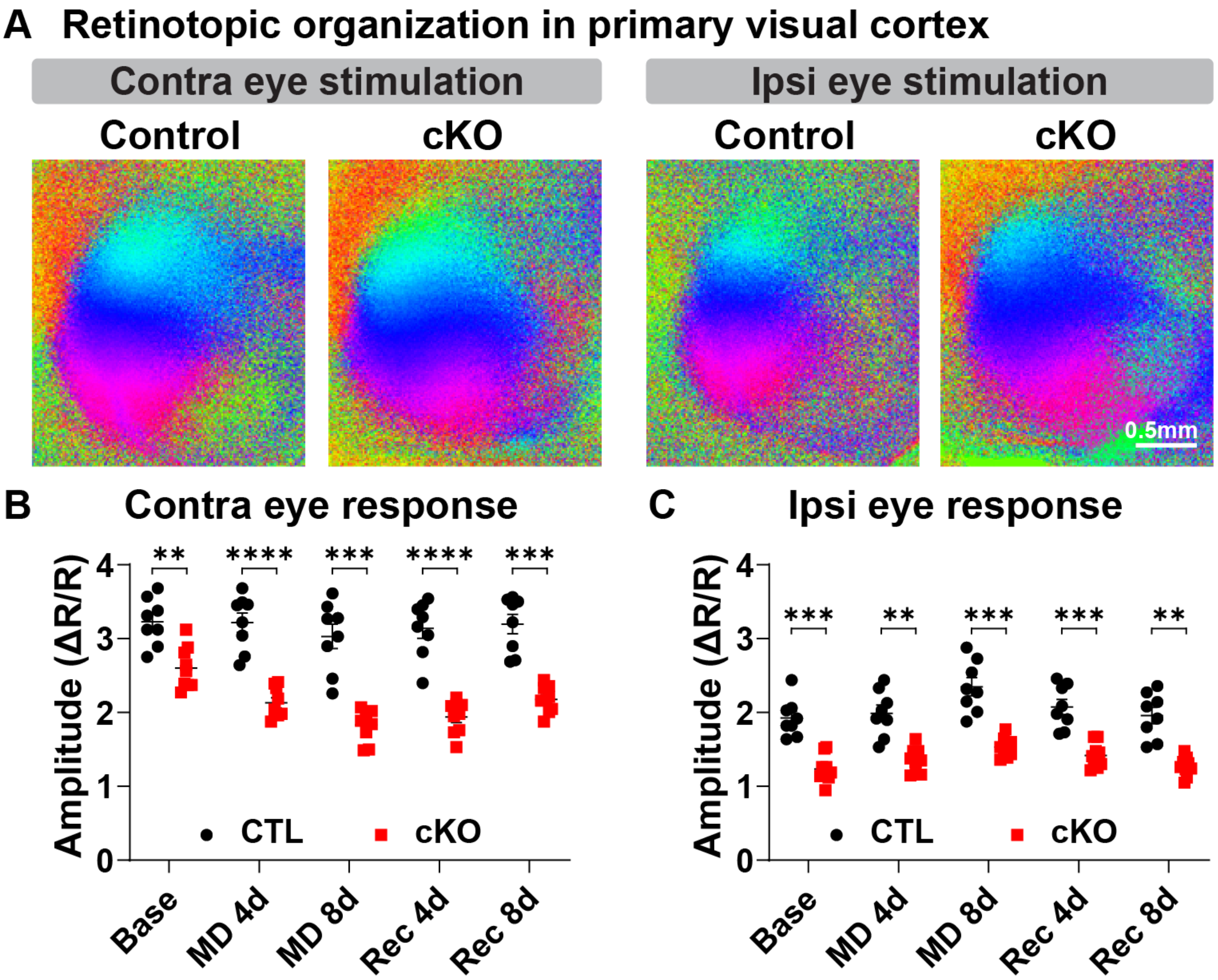


**Fig. S7.**

**Retinotopic organization and amplitude of visual cortex responses to visual stimulation in adult control and Myrf cKO mice.** (**A**) Example intrinsic signal images of retinotopy in primary visual cortex. (**B**, **C**) Amplitude of intrinsic signal imaging responses to stimulation of the contralateral deprived eye (contra) or ipsilateral non-deprived eye (ipsi) in binocular visual cortex of control (CTL) and cKO mice. Statistical details in Table S1. **$p<0.01$, ***$p<0.001$, ****$p<0.0001$.

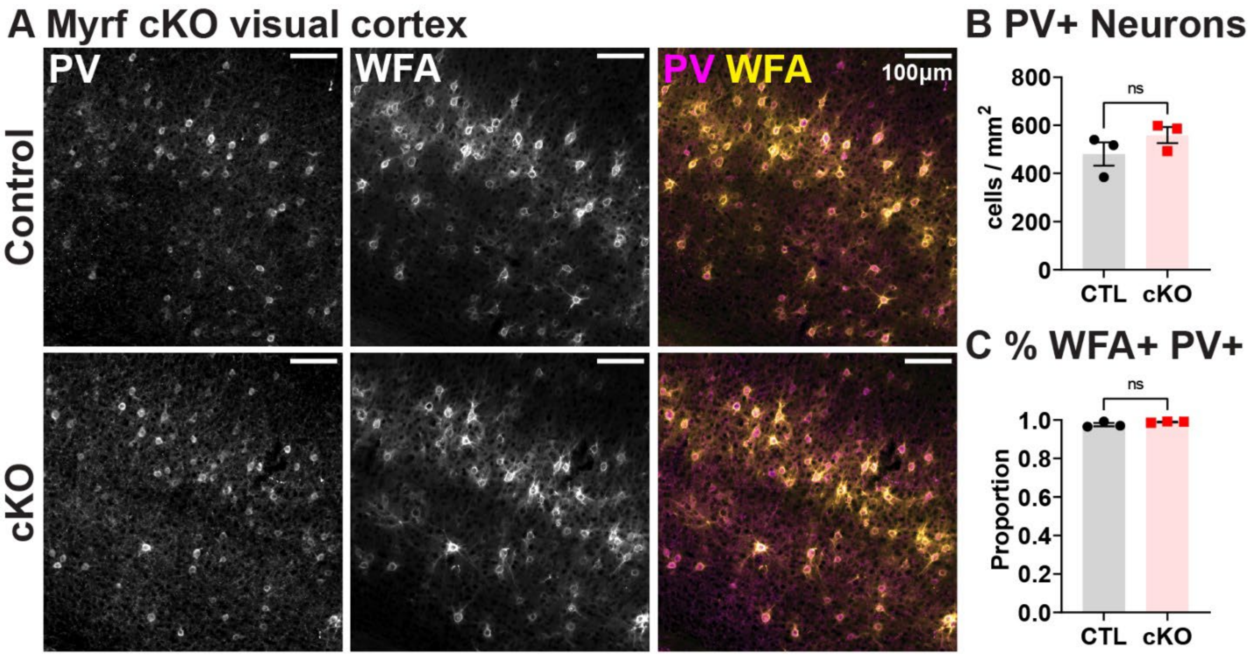


**Fig. S8.**

**Parvalbumin neuron density and perineuronal net coverage in adult control and Myrf cKO mice.** (**A**) Example images and quantification (**B**, **C**) of immunostaining for parvalbumin (PV) and perineuronal nets (WFA) in adult visual cortex of control (CTL) and cKO mice. Statistical details in Table S1. ns = not significant.

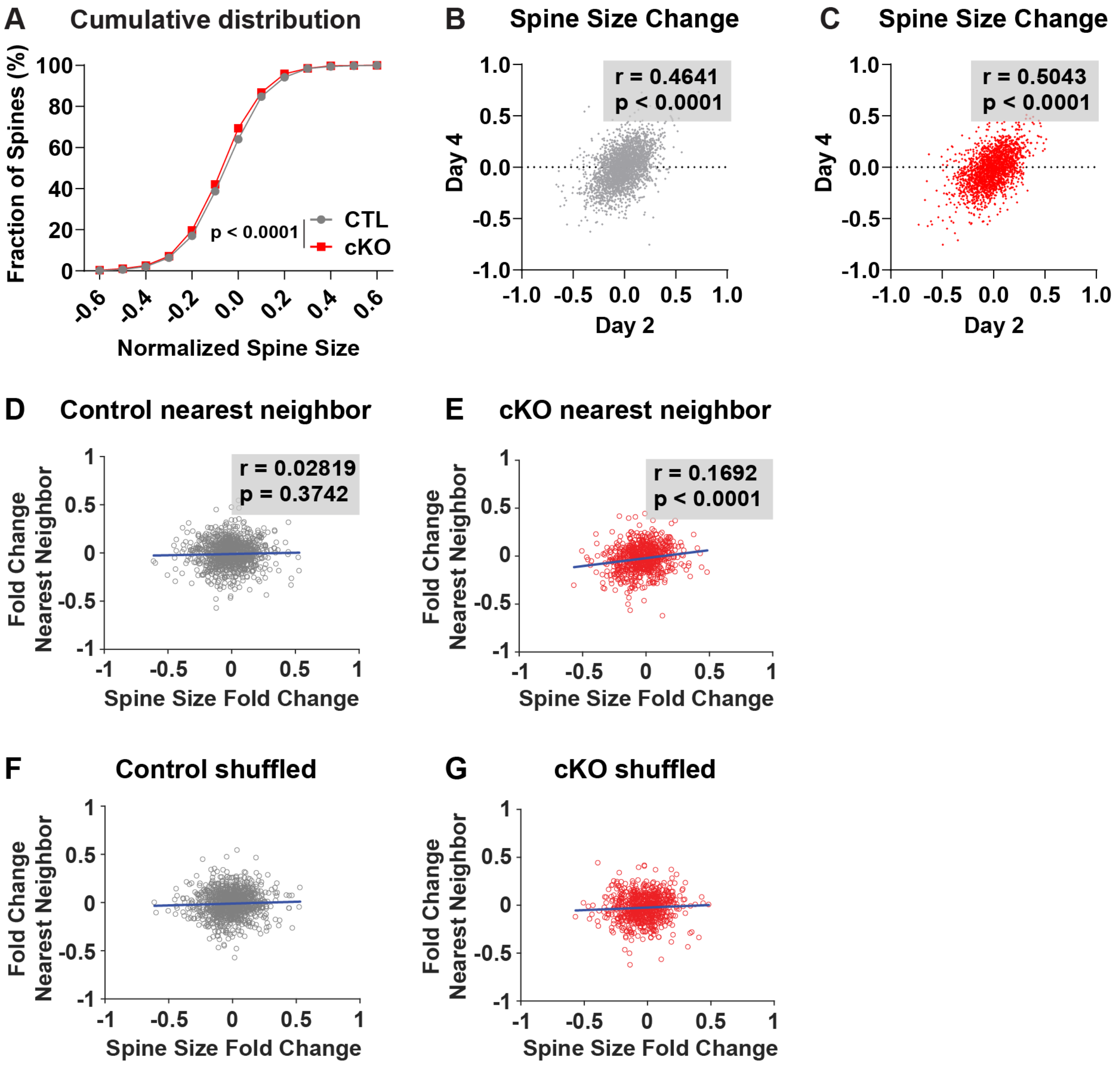


**Fig. S9.**

**Spine size changes following monocular deprivation in adult control and Myrf cKO mice.** (**A**) Cumulative distribution plot of spine size changes in control (CTL) and cKO mice after four days of monocular deprivation. (**B**, **C**) Correlation of spine size changes after two days of monocular deprivation with spine size changes after four days of monocular deprivation in control and cKO mice. (**D**, **E**) Correlation of average size change following monocular deprivation for a given spine and size change of its nearest neighbor in control and cKO mice. (**F**, **G**) Example correlation of nearest neighbor spine changes in one set of shuffled spine pairings for control and cKO mice. Statistical details in Table S1.

**Table S1.**
**Statistical analysis**

| **Figure** | **Sample size** | **Statistical test** | **Values** |
|---|---|---|---|
| 1H | 8 mice | Paired t-test (two-tailed) | t = 5.494, p = 0.0009 |
| 1I | 8 mice | Paired t-test (two-tailed) | t = 3.06, p = 0.0183 |
| 1J | 8 mice | Paired t-test (two-tailed) | t = 0.3199, p = 0.7584 |
| 1K | 8 mice | Paired t-test (two-tailed) | t = 3.84, p = 0.0064 |
| 1L | 5 mice | Paired t-test (two-tailed) | t = 3.867, p = 0.018 |
| 1M | 5 mice | Paired t-test (two-tailed) | t = 1.64, p = 0.1763 |
| 2B | 3-6 mice per age, per genotype | Two-way ANOVA followed by Sidak's multiple comparisons test | Genotype:<br>F = 165.9, p < 0.0001<br>Interaction:<br>F = 9.55, p = 0.0002<br><br>CTL – cKO<br>P28: p = 0.1808<br>P35: p < 0.0001<br>P45: p < 0.0001<br>P60: p < 0.0001 |
| 2C | 3-4 mice per age, per genotype | Two-way ANOVA followed by Sidak's multiple comparisons test | Genotype:<br>F = 0.7504, p = 0.7504<br>Interaction:<br>F = 0.0979, p = 0.7608<br><br>CTL vs cKO<br>P28: p > 0.9999<br>P60: p = 0.8672 |
| 2I | 8-9 mice per genotype | Two-way ANOVA followed by Sidak's multiple comparisons test | Genotype:<br>F = 30.64, p < 0.0001<br>Interaction:<br>F = 0.4264, p = 0.5237 |
| 2J | 8 mice | Two one-way repeated measures ANOVAs (one for contra and one for ipsi) followed by Tukey's multiple comparisons test | Contra<br>F = 1.733, p = 0.1804<br>Baseline vs MD4: p = 0.9863<br>Baseline vs MD8: p = 0.1922<br>MD4 vs MD8: p = 0.1814<br><br>Ipsi<br>F = 30.17, p < 0.0001<br>Baseline vs MD4: p = 0.5364<br>Baseline vs MD8: p = 0.0004<br>MD4 vs MD8: p = 0.0015 |
| 2K | 9 mice | Two one-way repeated measures ANOVAs (one for contra and one for ipsi) followed by Tukey's multiple comparisons test | Contra<br>F = 66.55, p < 0.0001<br>Baseline vs MD4: p = 0.0002<br>Baseline vs MD8: p < 0.0001<br>MD4 vs MD8: p = 0.0015<br><br>Ipsi<br>F = 20.66, p < 0.0001<br>Baseline vs MD4: p = 0.0575<br>Baseline vs MD8: p = 0.0013<br>MD4 vs MD8: p = 0.0053 |
| 2L | 8-9 mice per genotype | Unpaired t-test (two-tailed) | t = 5.082, p = 0.0001 |

| | | | |
|---|---|---|---|
| 2M | 8-9 mice per genotype | Two-way ANOVA followed by Sidak's multiple comparisons test | Genotype:<br>F = 36.22, p < 0.0001<br>Interaction:<br>F = 15.62, p < 0.0001<br><br>CTL vs cKO<br>MD 4d: p = 0.0003<br>MD 8d: p = 0.0007<br>Rec 4d: p = 0.0002<br>Rec 8d: p = 0.0013 |
| 2N | 8-9 mice per genotype | Two-way ANOVA followed by Sidak's multiple comparisons test | Genotype:<br>F = 1.247, p = 0.3008<br>Interaction:<br>F = 1.922, p = 0.1859<br><br>CTL vs cKO<br>MD 4d: p = 0.5115<br>MD 8d: p = 0.9725<br>Rec 4d: p = 0.2595<br>Rec 8d: p = 0.9961 |
| 3D | 10 control mice (111 dendrites)<br><br>10 cKO mice (97 dendrites) | Two-way repeated measures ANOVA followed by Holm-Sidak's multiple comparisons test | Genotype:<br>F = 7.394, p = 0.0071<br>Interaction:<br>F = 2.115, p = 0.0972<br><br>CTL vs cKO<br>Day -2: p = 0.0311<br>Day 0: p = 0.0103<br>Day 2: p = 0.0095<br>Day 4: p = 0.0391 |
| 3E | 10 control mice (111 dendrites)<br><br>10 cKO mice (97 dendrites) | Mixed-effects analysis (REML) followed by Holm-Sidak's multiple comparisons test | Genotype:<br>F = 8.82, p = 0.0033<br>Interaction:<br>F = 0.6344, p = 0.5931<br><br>CTL vs cKO<br>Day -2: p = 0.3037<br>Day 0: p = 0.0103<br>Day 2: p = 0.0178<br>Day 4: p = 0.0586 |
| 3F | 10 control mice (111 dendrites)<br><br>10 cKO mice (97 dendrites) | Mixed-effects analysis (REML) followed by Holm-Sidak's multiple comparisons test | Genotype:<br>F = 13.9, p = 0.0002<br>Interaction:<br>F = 1.101, p = 0.3484<br><br>CTL vs cKO<br>Day -2: p = 0.0288<br>Day 0: p = 0.0048<br>Day 2: p = 0.0521<br>Day 4: p = 0.192 |
| 3G | 10 control mice (111 dendrites)<br><br>10 cKO mice (97 dendrites) | Mixed-effects analysis (REML) followed by Holm-Sidak's multiple comparisons test | Genotype:<br>F = 3.483, p = 0.0634<br>Interaction:<br>F = 0.9613, p = 0.4107 |

| | | | |
|---|---|---|---|
| | | | CTL vs cKO<br>Day -2: p = 0.7148<br>Day 0: p = 0.2422<br>Day 2: p = 0.1721<br>Day 4: p = 0.1721 |
| 3H | 10 control mice (111 dendrites)<br><br>10 cKO mice (97 dendrites) | Two-way repeated measures ANOVA followed by Holm-Sidak's multiple comparisons test | Genotype:<br>F = 2.471, p = 0.1175<br>Interaction:<br>F = 2.64, p = 0.0486<br><br>CTL vs cKO<br>Day -2: p = 0.7333<br>Day 0: p = 0.7241<br>Day 2: p = 0.9743<br>Day 4: p = 0.0110 |
| 4C | 10 control mice (110 dendrites)<br><br>10 cKO mice (96 dendrites) | Unpaired t-test (two-tailed) | t = 0.3287, p = 0.7427 |
| 4D | 10 control mice (110 dendrites)<br><br>10 cKO mice (96 dendrites) | Unpaired t-test (two-tailed) | t = 2.826, p = 0.0052 |
| 4E | 10 control mice (110 dendrites)<br><br>10 cKO mice (96 dendrites) | Unpaired t-test (two-tailed) | t = 2.56, p = 0.0.0112 |
| 4F | 10 control mice (110 dendrites)<br><br>10 cKO mice (96 dendrites) | Unpaired t-test (two-tailed) | t = 1.627, p = 0.1052 |
| 4G | 10 control mice (996 spine pairs)<br>10 cKO mice (704 spine pairs) | Monte Carlo p value calculated by summing the tail of the histogram from 10000 shuffled spine pairings | p = 0.686 |
| 4H | 10 control mice (996 spine pairs)<br>10 cKO mice (704 spine pairs) | Monte Carlo p value calculated by summing the tail of the histogram from 10000 shuffled spine pairings | p = 0.939 |
| 4I | 10 control mice (996 spine pairs)<br>10 cKO mice (704 spine pairs) | Monte Carlo p value calculated by summing the tail of the histogram from 10000 shuffled spine pairings | p = 0.913 |
| 4J | 10 control mice (996 spine pairs)<br>10 cKO mice (704 spine pairs) | Monte Carlo p value calculated by summing the tail of the histogram from 10000 shuffled spine pairings | p = 0.66 |

| | | | |
|---|---|---|---|
| 4K | 10 control mice (996 spine pairs) 10 cKO mice (704 spine pairs) | Monte Carlo p value calculated by summing the tail of the histogram from 10000 shuffled spine pairings | $p = 0.862$ |
| 4L | 10 control mice (996 spine pairs) 10 cKO mice (704 spine pairs) | Monte Carlo p value calculated by summing the tail of the histogram from 10000 shuffled spine pairings | $p = 0.001$ |
| 4M | 10 control mice (996 spine pairs) 10 cKO mice (704 spine pairs) | Monte Carlo p value calculated by summing the tail of the histogram from 10000 shuffled spine pairings | $p = 0.002$ |
| 4N | 10 control mice (996 spine pairs) 10 cKO mice (704 spine pairs) | Monte Carlo p value calculated by summing the tail of the histogram from 10000 shuffled spine pairings | $p = 0.022$ |
| S1B | 8 mice | Paired t-test (two-tailed) | $t = 2.565$, $p = 0.0373$ |
| S1C | 8 mice | Paired t-test (two-tailed) | $t = 2.176$, $p = 0.066$ |
| S1F | 5 mice | Paired t-test (two-tailed) | $t = 3.637$, $p = 0.022$ |
| S3A | 4-6 mice per genotype | Unpaired t-test (two-tailed) | $t = 0.1984$, $p = 0.8477$ |
| S3B | 4-5 mice per genotype | Unpaired t-test (two-tailed) | $t = 3.461$, $p = 0.0105$ |
| S4B | 4-6 mice per genotype | Unpaired t-test (two-tailed) | $t = 2.982$, $p = 0.0176$ |
| S4C | 4-6 mice per genotype | Unpaired t-test (two-tailed) | $t = 2.195$, $p = 0.0594$ |
| S5A | 4 mice per genotype | Unpaired t-test (two-tailed) | $t = 0.3735$, $p = 0.7216$ |
| S5C | 4 mice per genotype | Unpaired t-test (two-tailed) | $t = 0.595$, $p = 0.5736$ |
| S5D | 4 mice per genotype | Unpaired t-test (two-tailed) | $t = 0.4586$, $p = 0.6627$ |
| S7B | 8-9 mice per genotype | Two-way ANOVA followed by Sidak's multiple comparisons test | Genotype:<br>$F = 55.3$, $p < 0.0001$<br>Interaction:<br>$F = 11.73$, $p < 0.0001$<br><br>CTL vs cKO<br>Base: $p = 0.0041$<br>MD 4d: $p < 0.0001$<br>MD 8d: $p = 0.0004$<br>Rec 4d: $p < 0.0001$<br>Rec 8d: $p = 0.0002$ |
| S7C | 8-9 mice per genotype | Two-way ANOVA followed by Sidak's multiple comparisons test | Genotype:<br>$F = 42.31$, $p < 0.0001$<br>Interaction:<br>$F = 2.183$, $p = 0.0816$<br><br>CTL vs cKO<br>Base: $p = 0.0002$ |

| | | | |
|---|---|---|---|
| | | | MD 4d: p = 0.0022<br>MD 8d: p = 0.0008<br>Rec 4d: p = 0.0009<br>Rec 8d: p = 0.0011 |
| S8B | 3 mice per genotype | Unpaired t-test (two-tailed) | t = 1.331, t = 0.2538 |
| S8C | 3 mice per genotype | Unpaired t-test (two-tailed) | t = 1.635, t = 0.1773 |
| S9A | 10 control mice (3484 spines)<br>10 cKO mice (2438 spines) | Kolmogorov-Smirnov test | p < 0.0001 |
| S9B | 10 control mice (3809 spines) | Pearson r correlation | r = 0.4641, p < 0.0001 |
| S9C | 10 cKO mice (3484 spines) | Pearson r correlation | r = 0.5043, p < 0.0001 |
| S9D | 10 control mice (996 spine pairs) | Pearson r correlation | r = 0.02819, p = 0.3742 |
| S9E | 10 cKO mice (704 spine pairs) | Pearson r correlation | r = 0.1692, p < 0.0001 |
| S9F, G | N/A | Example of one shuffled spine pairing as part of Monte Carlo simulation; no statistics involved | N/A |

**Table S2.**
**Primary antibodies**

| Antibody | Source | Identifier | Concentration |
|---|---|---|---|
| Rabbit anti-ASPA | GeneTex | Cat# GTX113389; RRID AB_2036283 | 1:1000 |
| Chicken anti-GFP | Rockland | Cat# 600-901-215; RRID AB_1537403 | 1:1000 |
| Rat anti-MBP | Millipore | Cat# MAB386; RRID AB_94975 | 1:200 |
| Rabbit anti-PDGFRα | W.B. Stallcup | N/A | 1:200 |
| Rabbit anti-cleaved Caspase3 | Cell Signaling | Cat# 9661S; RRID AB_2341188 | 1:200 |
| Mouse anti-GFAP | Millipore | Cat# MAB360; RRID AB_11212597 | 1:1000 |
| Goat anti-Sox9 | R&D Systems | Cat# AF3075; RRID AB_2194160 | 1:2000 |
| Rabbit anti-Iba1 | Wako | Cat# 019-19741; RRID AB_839504 | 1:1000 |
| Mouse anti-NF-L Degenotag | Encor | Cat# MCA-1D44; RRID AB_2923483 | 1:1000 |
| Rabbit anti-NF-H | Abcam | Cat# ab8135; RRID AB_306298 | 1:1000 |
| Mouse anti-PV | Swant | Cat# 235; RRID AB_10000343 | 1:1000 |
| Biotinylated WFA | Vector Labs | Cat# B-1355; RRID AB_2336874 | 1:400 |
